# First genome assembly of the order Strepsiptera using PacBio HiFi reads reveals a miniature genome

**DOI:** 10.1101/2024.05.07.592988

**Authors:** María Isabel Castaño, Xinhai Ye, Floria MK Uy

**Affiliations:** Department of Biology, University of Rochester, Rochester, NY, USA; College of Advanced Agriculture Science, Zhejiang A&F University, Hangzhou, Zhejiang, China

**Keywords:** Genome, HiFi reads, parasite, sexual dimorphism, Strepsiptera, twisted-wing insect, *Xenos peckii*

## Abstract

Twisted-wing insects (Strepsiptera) are an enigmatic order of parasites with unusual life histories and striking sexual dimorphism. Males emerge from hosts as free-living winged adults, while females from most species remain as endoparasites that retain larval traits. Due to scarce genomic data and phylogenetic controversies, Strepsiptera was only recently placed as the closest living relative to beetles (Coleoptera). Here, we report the first PacBio HiFi genome assembly of the strepsipteran *Xenos peckii* (Xenidae). This *de novo* assembly size is 72.1 Mb, with a BUSCO score of 87.4%, N50 of 7.3 Mb, 23.4% GC content and 38.41% repeat content. We identified 8 contigs that contain > 75% of the assembly and reflect the haploid chromosome number reported from karyotypic data, and 3 contigs that exhibit sex chromosome coverage patterns. Additionally, the mitochondrial genome is 16,111 bp long and has 34 genes. This long-read assembly for Strepsiptera reveals a miniature genome and provides a unique tool to understand complex genome evolution associated with a parasitic lifestyle and extreme sexual dimorphism.

## Background & Summary

The order Strepsiptera, commonly known as twisted-wing insects, are a group of enigmatic obligate parasites, with 9 extant families and approximately 600 described species^1^. High diversification rates associated with adaptive specialization to their hosts has allowed them to parasitize 7 orders and 34 families of insects^2,3^. More than 97% of Strepsiptera species are characterized by extreme sexual dimorphism, except for the ancestral family Mengenillidae where winged males and apterous females emerge as free-living adults^3,4^. In the remaining families, males pupate and emerge as free-living winged adults, while adult neotenic females lack a pupal instar, and remain as obligate endoparasites inside their host^5^. The grub-like females lack a distinctive head, thorax, abdomen or body appendages and are viviparous, with some of the highest fecundity rates reported for any insect ^6^.

A parasitic lifestyle coupled with unusual life history traits have been linked to unique genetic characteristics: exceptionally small genomes reported by flow cytometry, different levels of endoreduplication between males and females, and high rates of molecular evolution^6,7^. These unusual genetic and morphological features, combined with scarce genomic data available for the order, hindered efforts to place this order in the insect phylogeny for decades, calling it the Strepsiptera problem^8^. However, recent studies used a whole genome shotgun sequencing approach of the basal species *Mengenilla moldrzyki* (Mengenillidae) to resolve the enigma and placed Strepsiptera confidently as the closest living relative to beetles (Coleoptera)^9,10^. The only other genomic data available for these unique parasites is based off short read whole genome sequencing and shows intriguing evidence for partial dosage compensation in the derived species, *Xenos vesparum* (Xenidae)^11^.

Parasites often use metabolic resources from their hosts^12^. Therefore, selection for metabolic efficiency, such as costs of replicating DNA, can result in smaller genomes^13,14^. Similarly, organisms that undergo massive changes of their body plan when transitioning to an obligate endoparasitic life, often experience extensive gene losses as a tradeoff ^15,16^. In Strepsiptera, genome size reduction or potential gene losses associated with host specialization and extreme sexual dimorphism combine compelling evolutionary scenarios to understand genome evolution^6^. In fact, the only cytogenetic data available for Strepsiptera indicates they have XY sex chromosomes and substantial variation in diploid chromosome number within the same family (i.e., 16 chromosomes in *Xenos peckii* and 8 chromosomes reported for *Brasixenos sp.* from Brazil: Xenidae: Strepsiptera)^17^. Xenidae is one of the most derived families in Strepsiptera and exhibits a high degree of host specialization. The ability to colonize new host lineages resulted in a constant net diversification rate that has produced between 77 and 96 potentially cryptic species^18^.

Throughout the world, many species of *Xenos* consistently infect social wasps of the genus *Polistes* (Figure 1)^19,20^. Studies in two parallel *Xenos*-*Polistes* systems in Europe and North America reveal consistent behavioral and physiological manipulation strategies ^5,21,22^. Parasitic effects include hindering host reproduction and manipulating wasps to abandon their colony to form aggregations that facilitate mating of adult parasites. Thus, members of *Xenos* evolve in a continuous arms race with their hosts to overcome the host immune system^23^, which also may influence genome evolution. As the genomic revolution advances, sequencing of high-quality Strepsiptera genomes is key to understanding the molecular mechanisms underpinning host manipulation as well as the forces driving genome size evolution in this enigmatic group of insects.

**Figure 1.**
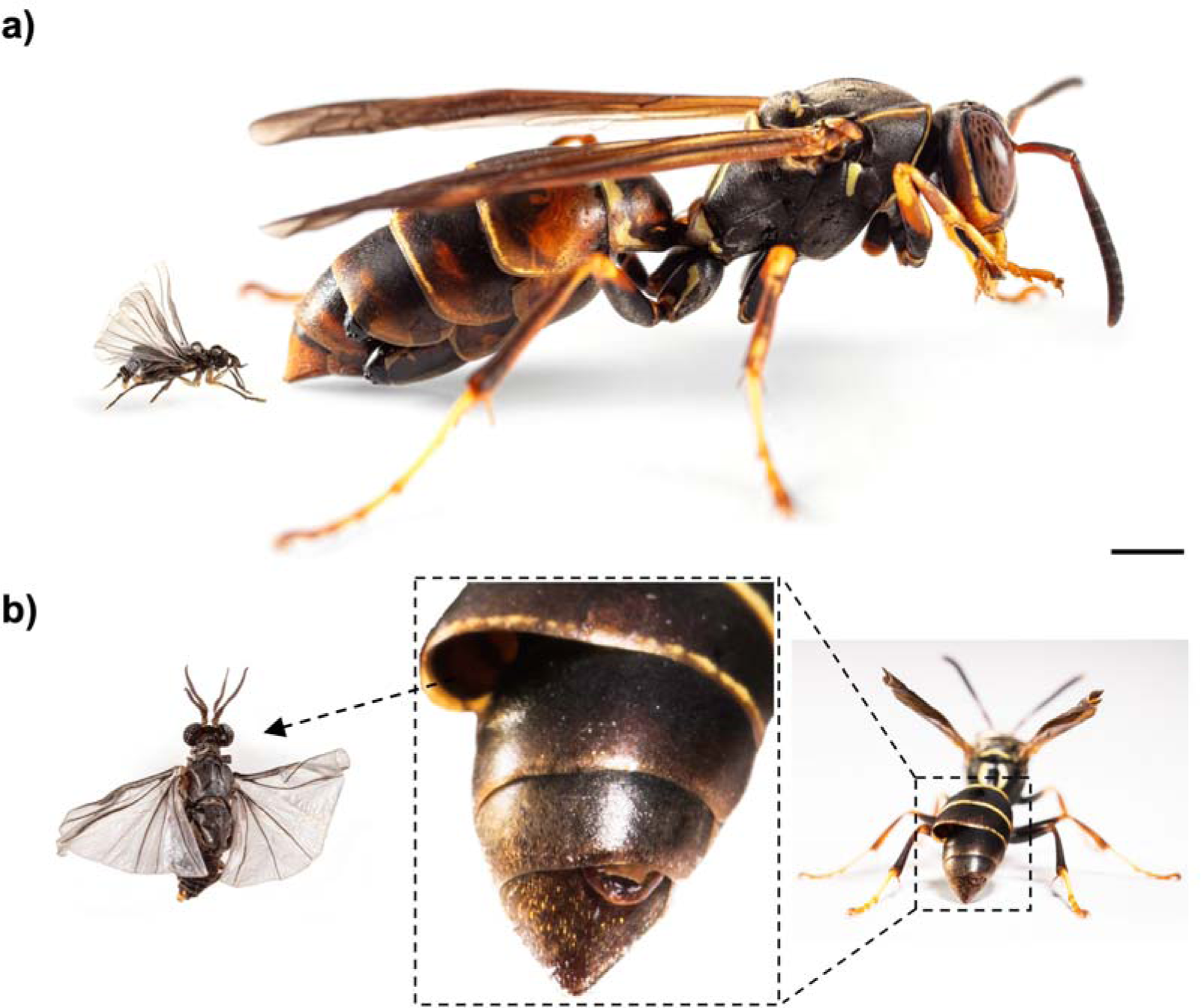
Description of the study system. **A**) A paper wasp, *Polistes fuscatus,* infected with two male parasites lodged in the abdominal cavity (right). An adult, free-living male *Xenos peckii* emerged from this host (left) (Scale Bar = 2 mm). **B)** Detailed morphology of males and females. Left: A free-living adult male that emerges from the abdomen of the host. Center: close-up of wasp abdomen that shows the protruding cephalothorax of an obligate endoparasitic female between abdominal segments and the empty pupal case where the male parasite emerged from. Females have a flattened dorsoventrally flattened cephalothorax and lack all adult insect features (i.e., eyes, antennae, mouth parts, legs, wings). Right: infected wasp host and size of parasites lodged in its abdomen for scale.

In this study, we present the first *de novo* genome assembly of the order Strepsiptera – A male *Xenos peckii* (Strepsiptera: Xenidae) – using PacBio HiFi sequencing technology. After filtering out bacterial scaffolds, the haploid assembly size is 72.1 Mb, assembled in 227 contigs, a contig N50 of 7.3 Mb, a GC content of 23.4%, BUSCO score of 87.4% and a repetitive element content of 38.41%. This is the most complete and contiguous genome assembly available for the order Strepsiptera up to date. More than 75% of the genome is contained in 8 major contigs (L75=8) and the longest contig is 8.76Mb. The 8 major contigs reflect the haploid number of chromosomes reported from karyotypic data (n=8)^17^. Furthermore, we identified 3 contigs that exhibit sex chromosome type coverage patterns (half depth of coverage in males). Additionally, we assembled and annotated the mitochondrial genome, which is 16111bp long and has 34 genes including transfer RNAs. The size and gene content of the mitochondrial genome is similar to that of published mitogenomes for *Xenos vesparum* found in Europe and its close relative, *Xenos yangi* from China^24,25^. Recent work has resolved that *Xenos cf. moutoni* and X. yangi are the same species due to a reported genetic divergence between 0 and 0.14. To our knowledge, the twisted-wing parasite genome we report is among the 10 smallest insect genomes reported in the literature, and the smallest insect genome assembled using long PacBio HiFi reads^26^. Our high-quality genomic data provides a unique tool to improve our understanding of the complex genome evolution associated with extreme sexual dimorphism coupled with a parasitic lifestyle. Additionally, this genome will facilitate the exploration of the molecular mechanisms underlying host and twisted-wing parasite coevolution.

## Methods

### Specimen collection and sample preparation

During June of 2021, we used entomological nets to collect infected social wasps, *Polistes fuscatus*, at the Robert H. Treman State Park located in Ithaca, NY, USA. Infection and stage of development of *X. peckii* was confirmed by the extrusions in the host abdomen, which are morphologically distinct for developing female and male parasites^5^. We transported the infected wasps to the University of Rochester in disposable transparent deli cups that are pre-punched to increase air flow. Each infected wasp was housed individually in a deli cup with a water vial and a sugar cube and kept under controlled conditions that included: temperature of 25°C, 40% humidity, and full spectrum light for 12 hours. We inspected infected wasps daily during the extrusion period of 3-7 days and 9-17 days for male and female parasites, respectively. After emerging from a host, male parasites were frozen with dry ice and stored at −80°C. A single female was dissected out of its host and preserved the same way. A single male *X. peckii* was preserved for an ultra-low DNA input sample preparation and subsequent long-read sequencing, while an additional male and the female were preserved for short-read sequencing. Finally, we froze one more single male in liquid nitrogen for subsequent extraction of RNA, sequencing, and genome annotation.

### DNA and RNA extraction, library preparation, sequencing and pre-assembly read processing

Library preparation and sequencing for PacBio HiFi data was performed at the DNA sequencing and Genotyping Center of the Delaware Biotechnology Institute at University of Delaware. Total genomic DNA was isolated using the Qiagen Megattract kit (QIAGEN, shanghai, China) following the manufacturer’s instructions. One SMRTbell library of circular consensus sequencing (CCS) was constructed according to the Low DNA input PacBio protocol with additional bead cleaning. The library was then sequenced on one PacBio Sequel II SMRT cell platform in CCS mode, which calls consensus reads from subreads that are generated after multiple passes of the enzyme around the circularized template (Pacific Biosciences). Sequencing yielded 8.6 Gb of data and ∼0.9 million HiFi (high-fidelity) reads from the male *X. peckii* sample (Table 1).

**Table 1.**
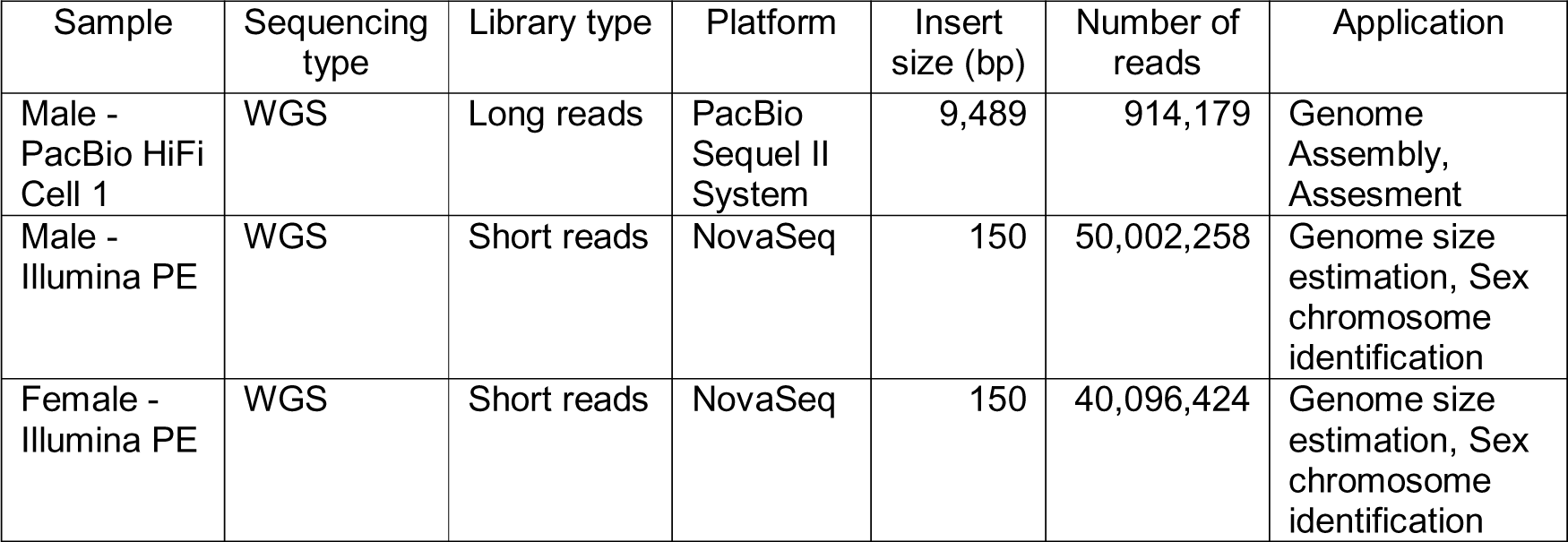
General statistics of raw sequencing reads used for *X. peckii* genome assembly and genome size estimation.

| Sample | Sequencing type | Library type | Platform | Insert size (bp) | Number of reads | Application |
| --- | --- | --- | --- | --- | --- | --- |
| Male - PacBio HiFi Cell 1 | WGS | Long reads | PacBio Sequel II System | 9,489 | 914,179 | Genome Assembly, Assesment |
| Male - Illumina PE | WGS | Short reads | NovaSeq | 150 | 50,002,258 | Genome size estimation, Sex chromosome identification |
| Female - Illumina PE | WGS | Short reads | NovaSeq | 150 | 40,096,424 | Genome size estimation, Sex chromosome identification |

**Table 2.**
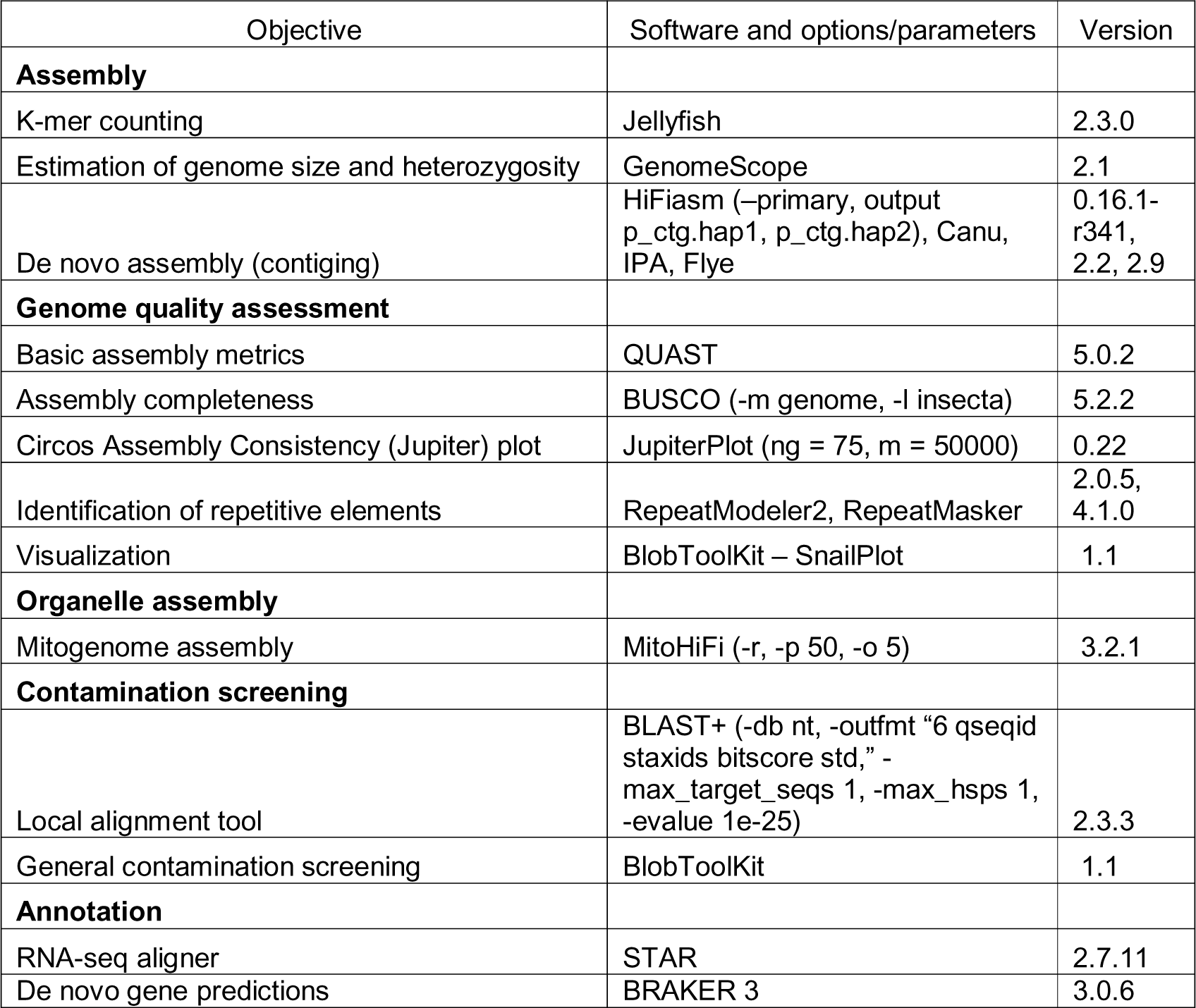
Software and version used for analyses in this study with corresponding parameters, if different from the default options.

We extracted Genomic DNA from the additional male and the female *X. peckii* using a DNeasy Blood and Tissue kit (QIAGEN, Valencia, CA, USA) for library preparation and short read whole genome Illumina sequencing. RNA from a whole male was extracted with the RNeasy Mini Kit, implementing a DNase digestion step with the RNase-Free DNase Set (QIAGEN, Valencia, CA, USA). DNA and RNA samples were received and validated via standard quality control procedures at Novogene Co., Sacramento, CA. To construct the genomic DNA library, DNA was sheared into short fragments which were end repaired, A-tailed and ligated with Illumina adapter. Fragments with adapters were PCR amplified, size selected and purified. Each library was checked with Qubit and real time PCR for quantification, and then pooled for sequencing on a NovaSeq PE 150 Illumina platform (Novogene Co. Ltd. Sacramento, CA). Sequencing results generated 7.5Gb and 6.0Gb of data for the male and female samples, respectively (Table 1). After sequencing, we assessed read quality using FastQC^27^. Then, we filtered reads for quality and length, and trimmed adapter sequences and PolyG tails using fastp^28^ with the options −l 40. Then we re-assessed the quality of the processed reads, which we used to validate the genome size, heterozygosity and repetitive content estimations and to identify sex chromosomes based on coverage patterns. The transcriptome data (total RNAseq) was also sequenced with the NovaSeq PE 150 platform, using an mRNA library preparation with standard poly A enrichment, which generated 6 GB of raw data to implement in the genome annotation.

### Genome assembly and size estimation

Prior to assembly, estimating features like genome size, rate of heterozygosity, and repetitive element content proves useful to inform the parameter values that should be used in subsequent steps. We used Jellyfish (v2.3.0)^29^ to count and compute a histogram of k-mer frequencies from the raw PacBio HiFi reads using the count (-C - m 21) and histo (–h 1,000,000) modules. Then we used the Jellyfish histogram output to run the online web tool of GenomeScope2 (http://qb.cshl.edu/genomescope/genomescope2.0) with the following parameters: K-mer length = 21, ploidy = 2 and max kmer coverage = 1,000,000^30^. The model suggests that *X. peckii* has an estimated genome size of 63,567,278 bp with 1.19% heterozygosity levels, and 36.4% of the genome composed of repetitive elements (Figure 2). The *X. peckii* genome we report here is one of the smallest insect genomes documented up to date^6^ (Table 3; See technical validation below).

**Figure 2.**
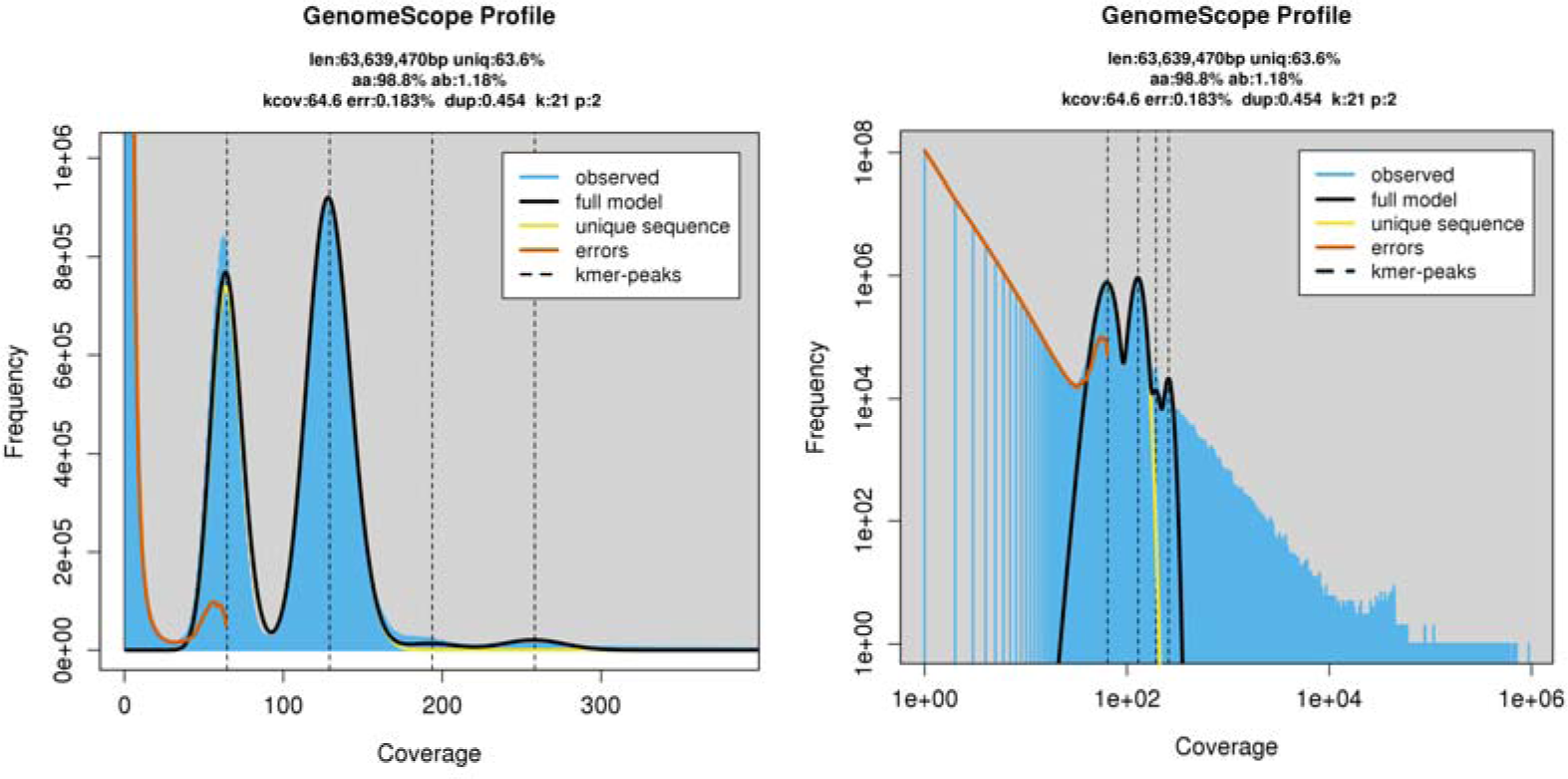
Genome size and content estimation using GenomeScope 2.0. Left: Genomescope2 k-mer (21) distribution from the adapter trimmed PacBio HiFi reads. K-mer depth is plotted against k-mer frequency for a given depth. The plot displays estimation of genome size (len), percentage of the genome that is not in repetitive elements (uniq), homozygous rate (aa), heterozygous rate (ab) mean k-mer coverage for heterozygous bases (kcov), read error rate (err), average rate of read duplications (dup), k-mer size used on the run (k) and ploidy (p). Left: Expected distribution according to the model. Right: Obtained distribution according to the model. The four peaks correspond to the mean coverage levels of the unique heterozygous, unique homozygous, repetitive heterozygous and repetitive homozygous sequences, respectively.

**Table 3.** GenomeScope estimation of genome structure using reads from different sequencing technologies. *X. cf. moutoni* and *X. yangi* are now considered the same species based on COI pairwise distances, as highlighted with an asterisk (Dong et al. 2022)

| Species | Type of reads | Genome size (bp) | Genome in repeats | Heterozygosity | Error rate | Duplications |
| --- | --- | --- | --- | --- | --- | --- |
| *X. peckii* (Male) | PacBio HiFi | 63,567,278 | 36.4% | 1.18% | 0.18% | 0.45% |
| *X. peckii* (Male) | Illumina PE 150bp | 67,923,714 | 43.7% | 1.1% | 0.25% | 1.97% |
| *X. peckii* (Female) | Illumina PE 150bp | 75,093,981 | 41.4% | 0.76% | 0.59% | 1.62% |
| *X. vesparum* (Female) | Illumina | 120,869,754 | 42.1% | 0.91% | 0.26% | 1.81% |
| *X. cf. moutoni* (Female)* | Illumina | 58,068,699 | 27.5% | 0.69% | 0.14% | 1.60% |

After a preliminary survey of the genome structure, we ran four different assembly software optimized for HiFi reads (i.e., Hifiasm^31^, Improved Phased Assembler IPA^32^, Hicanu^33^, Flye^34^) to gauge differences in assembly metrics. Hifiasm produced the highest quality genome (see technical validation below; Supplementary Table 1; Supplementary Figure 1). We assembled the HiFi reads without additional data into a draft haplotype-resolved genome assembly using Hifiasm v0.16.1-R341^31^. We then used the default parameters of Hifiasm to generate a primary and alternate assembly graph after aggressive purging of haplotypic duplications (-primary, −l 3) and three rounds of error correction^31^. The primary assembly produced by Hifiasm was 73,541,611 bp long, assembled in 229 contigs with an N50 of 7.3 Mb and a GC content of 23.6%. Both the primary (73Mb) and alternate (64Mb) assemblies produced by Hifiasm were similar to the estimated genome size produced by GenomeScope2. However, the primary assembly was slightly larger than the GenomeScope2 estimations (Table 3; Figure 2). High heterozygosity can result in spurious duplications that increase the genome size because two alleles from the same loci are included in the primary assembly. We used *Jupiter Plot* (-n=50000, ng=75)^35^ to do a synteny plot and compare completeness between the primary and alternate assemblies produced by Hifiasm. The primary assembly produced by Hifiasm is more contiguous and that the alternate assembly doesn’t include some of the largest contigs in the primary assembly (potential sex-chromosomes based on coverage patterns; Supplementary Figure 6). Thus, the small discrepancies (<10Mb) between the GenomeScope genome size estimations and the final assembly size may be due to assembler limitations when sorting or deduplicating repetitive regions.

### Genome quality assessment

We used Blobtools2 from the BlobToolKit suite^36^ to screen for contamination. First, we performed a BLASTn^37^ search of our assembly against the general RefSeq blast -nt database using the following parameters: -outfmt ‘6 qseqid staxids bitscore std’ - max_target_seqs 1 -num_threads 12 -evalue 1e-6. Then we used the function blobtools –add to create a BlobDir database that included the blast output (hits file), the read coverage (bam file) from mapping the raw reads back to the assembly, and the previously computed BUSCO scores (Busco summary file). We implemented the BlobToolKit v.1.1.1^36^ online Viewer (blobtools host ‘pwd’) to create a Blobplot with the hits of our bacterial contamination scan, and their respective coverage (Supplementary Figure 4). We removed two contigs that matched the phylum Proteobacteria. Specifically, one whole contig was the genome of *Wolbachia pipitensis* (see technical validation below; Supplementary Figure 5). Then, we used QUAST v3^38^ to assess the metrics of the curated assembly. The final assembly size was 72,105,243 Mb, assembled in 227 contigs with a GC content of 23.4%. We used the BlobtoolKit v.1.1.1^36^ to create a SnailPlot to visualize our assembly statistics (Table 4; Figure 3). This Whole Genome PacBio HiFi assembly project has been deposited at DDBJ/ENA/GenBank under the accession JAWUEG000000000.

**Figure 3.**
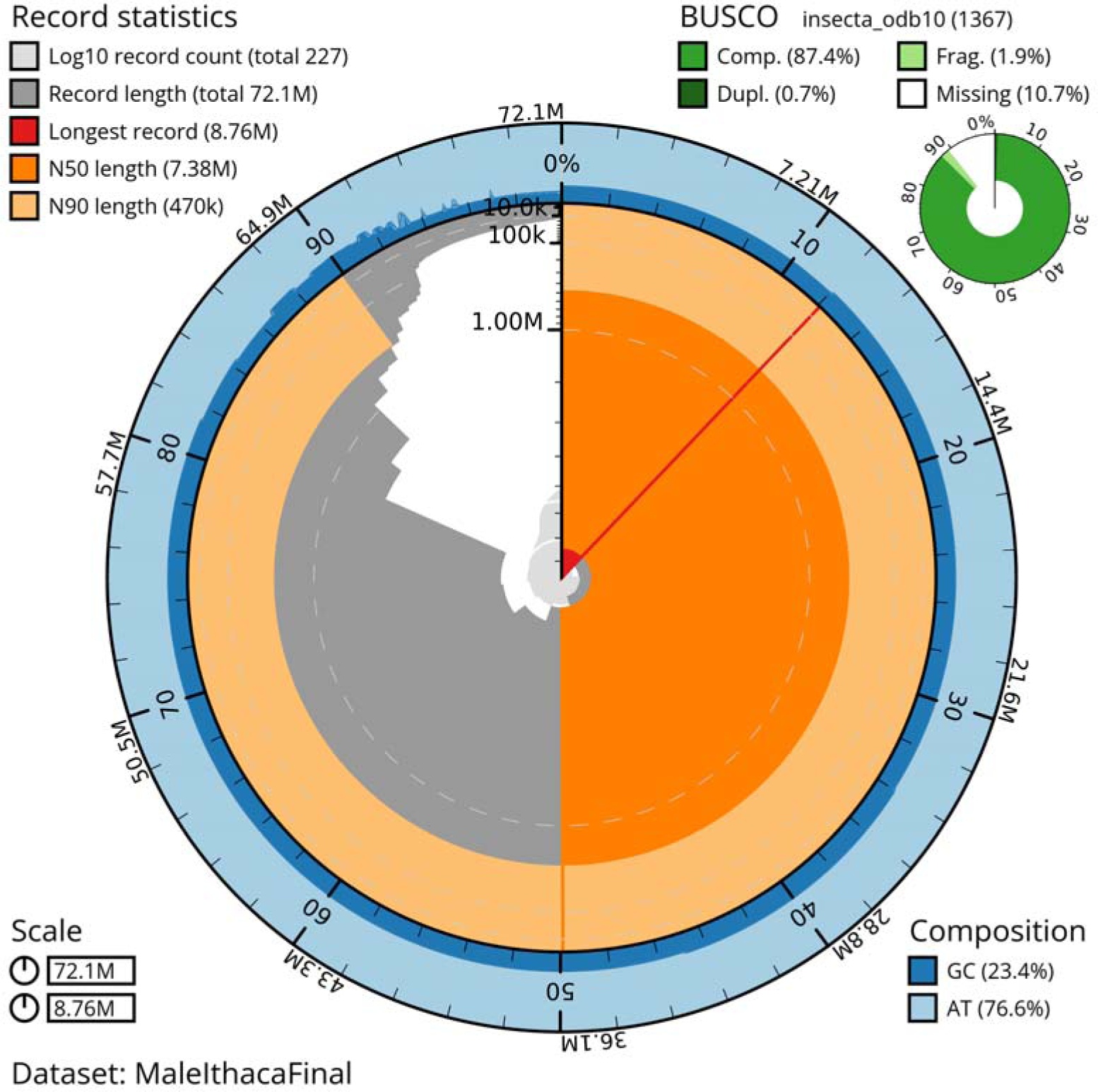
Snail plot assembly visualization using BlobTools 2.0. The contiguity and completeness of the *X. peckii* genome assembly after contamination screening is plotted as a circle that represents the full length of the assembly (∼72.1 Mb), assembled in 227 contigs. The N50 (7.38 Mb) is highlighted in dark orange and the N90 (470k) in light orange. The longest contig was 8.76 Mb (highlighted in red). The assembly has a uniform GC content of 23.4% and few contigs <100 Kb in length. The BUSCO scores are shown in the top right corner in green.

**Table 4.** Statistics produced by QUAST for final assembly using Hifiasm after removing bacterial contamination. All statistics are based on contigs of size >= 500 bp, unless otherwise noted (e.g., “# contigs (>= 0 bp)” and “Total length (>= 0 bp)” include all contigs).

|  |  |
| --- | --- |
| #Contigs ($\geq 0$ bp) | 227 |
| #Contigs ($\geq 1000$ bp) | 227 |
| #Contigs ($\geq 5000$ bp) | 227 |
| #Contigs ($\geq 10000$ bp) | 226 |
| #Contigs ($\geq 25000$ bp) | 111 |
| #Contigs ($\geq 50000$ bp) | 24 |
| Total length ($\geq 0$ bp) | 72105243 |
| Total length ($\geq 1000$ bp) | 72105243 |
| Total length ($\geq 5000$ bp) | 72105243 |
| Total length ($\geq 10000$ bp) | 72097022 |
| Total length ($\geq 25000$ bp) | 69922735 |
| Total length ($\geq 50000$ bp) | 67172384 |
| # contigs | 227 |
| Largest contig | 8758233 |
| Total length | 72105243 |
| GC (%) | 23.39 |
| N50 | 7378620 |
| N75 | 6101180 |
| L50 | 5 |
| L75 | 8 |
| #N's per 100 Kbp | 0 |

We evaluated assembly quality and completeness using Benchmarking Universal Single-copy Orthologs (BUSCO) v.5.2.2 ^39^ with the insecta_odb10 database. From a total of 1367 BUSCO groups searched, our assembly has 87.4 % of genes complete and present in single copy, 0.7% duplicated genes, 1.9% fragmented genes, and 10.4% of the genes missing (Figure 3). The underlying cause behind the high proportion of missing genes in our assembly will be clarified once more genomic data for the order becomes available.

### Mitochondrial genome assembly

We assembled the mitochondrial (mtDNA) genome of *X. peckii* from the raw PacBio HiFi reads with the MitoHiFi pipeline^40,41^, which uses a reference-guided method to perform the assembly. First, the pipeline implemented the software MitoFinder^41^ to scan all the complete mtDNA assemblies available on NCBI and downloads the reference mitogenome for the most closely related taxa. PacBio HiFi reads were mapped to the reference mitogenome using Minimap2 v.2.17^42^, and reads that were larger than the size of the reference were filtered out. The remaining filtered and mapped reads were assembled de novo with Hifiasm^31^ into a mitogenome. Finally, the MitoHifi pipeline annotates the mitochondria and calculates the depth of coverage along the sequence. Mitochondrial genomes are available for two *Xenos* species: *X. vesparum* (partial genome: 14,519bp NCBI Accession number DQ364229), *X yangi*, previously reported as *X. cf. moutoni, (*partial genome: 16,717bp NCBI Accession number MW222190) and *X. yangi* (complete genome: 15,324bp NCBI Accession number NC_067052.1)^24,25,43^. Following the MitoFinder output, we used the mitochondrial genome of *Xenos yangi,* as the starting reference sequence, because it is the only complete sequence available^43^. Then, we used MitoHiFi (-p 90 -o 5) to assemble the mitogenome. From a total of 30,623 HiFi reads that mapped to the reference mitogenome, we used only 28,091 reads that were shorter or equal to the reference *X. yangi* mitogenome (<15,324bp). The final mitogenome genome size assembled for *X. peckii* was 16,111bp and contains 34 genes including unique transfer RNAs (NCBI Reference Sequence: JAWUEG000000000; Supplementary Figure 3). We used BLAST+ v2.1^37^ to search for matches of our mitochondrial assembly in the nuclear genome and filtered out contigs from the nuclear genome with a percentage of sequence identity >99% and of smaller size than the mtDNA sequence ^44^.

### Mapping rate and coverage for X-linked contigs discovery

Cytogenetic data for the order Strepsiptera is only available for two species, and it indicates that they have heteromorphic X and Y chromosomes^11,45^. Specifically, in *Xenos peckii*, previously described as *Acroschismus wheeleri*, the diploid number of chromosomes (2n) was identified as 16 ^17^. To evaluate mapping rate and depth of coverage along the assembly we used Minimap2 v.2.17^42^ to map the raw long PacBio HiFi reads, as well as the short Illumina (150bp) PE reads, back to the final assembly. The male *X. peckii* HiFi reads had an average mapping rate of 99.43% and 97.8% coverage, with a mean depth of coverage of 117X. For the Illumina PE reads the mapping rate was 94.95% for the male and 77.61% for the female, with a mean depth of coverage of 97.27X and 63.73X respectively. We used the CIRCOS^46^ tool to visualize the depth of coverage through the first 11 contigs that contain > 90% of the genome. We identified three X-linked contigs (ctg02, ctg23 and ctg15), which presented a male to female coverage ratio of 0.5. Interestingly, ctg02 is also the largest contig in our assembly (Figure 4).

**Figure 4.**
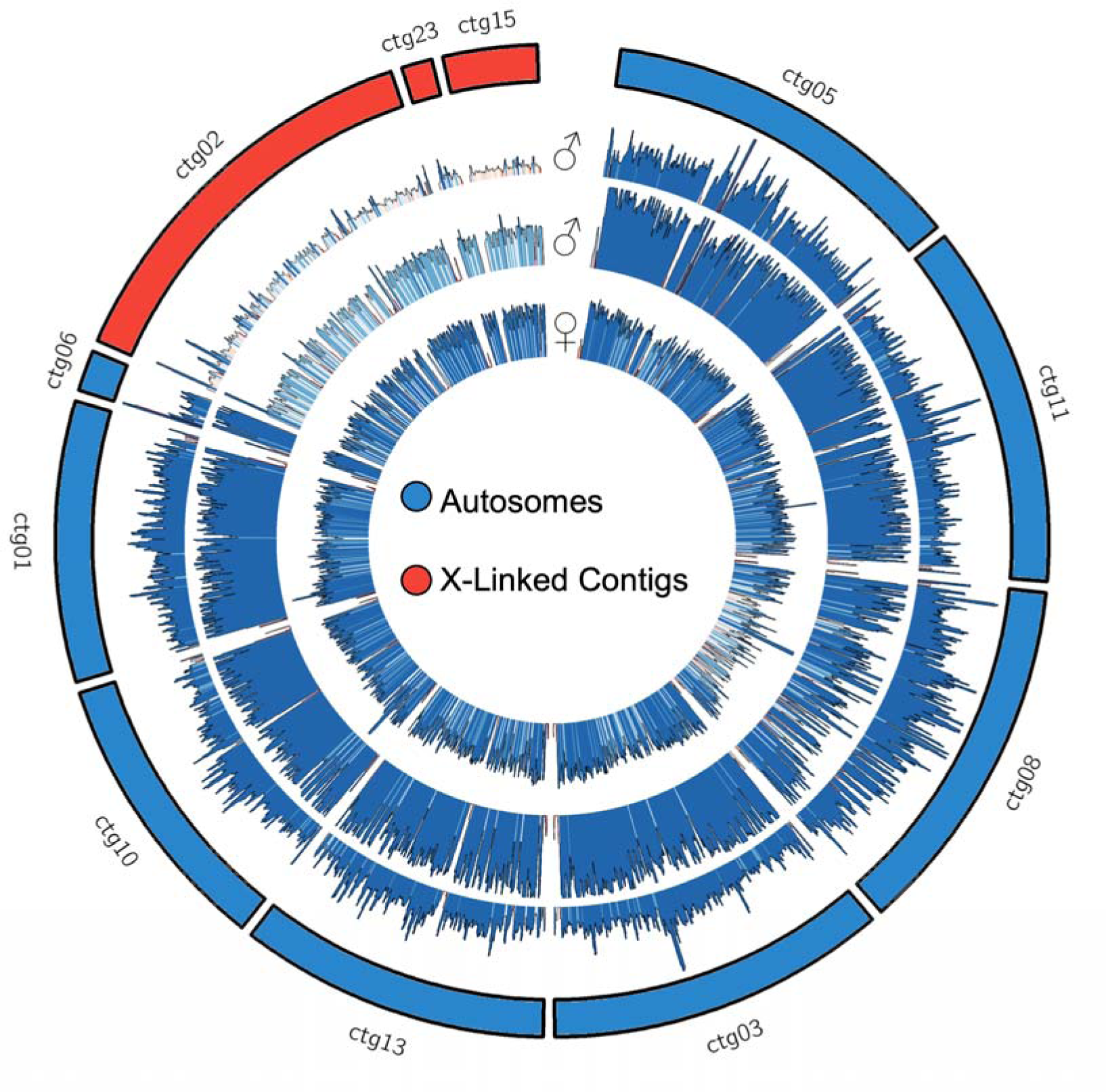
X-linked contig discovery based on depth of coverage. Average coverage over 50 Kb windows with 20Kb step size in the 11 largest contigs, that contain >90% of the genome. Three individual samples were used: the first external coverage plot corresponds to the HiFi reads of the *X. peckii* male used for the genome assembly. The middle coverage plot corresponds to short 150bp Illumina reads from another male, and the internal coverage plot corresponds short 150bp Illumina reads from a female. Potential X-linked contigs (red) have half the coverage in both male samples than the rest of the autosomes (blue). In contrast, the female sample has a uniform depth of coverage throughout the plot.

### Repeat Modeler

As this is the first high quality genome assembly of the order Strepsiptera and there are no repeat libraries available for the order, we identified and classified repetitive elements *de novo* using RepeatModeler v.2.0.133^47^. Then, we used RepeatMasker v.4.0.734^48^ to mask the genome assembly using the *de novo* library produced by RepeatModeler2. RepeatMasker masked 38.41% of the genome, a similar value to the repeat content suggested by GenomeScope2 before removing the bacterial contigs. Most of the repeats in the genome are unclassified (27.8%). LTR elements are the most abundant elements among the classified repeats (5.74%; Figure 5; Table 5).

**Figure 5.**
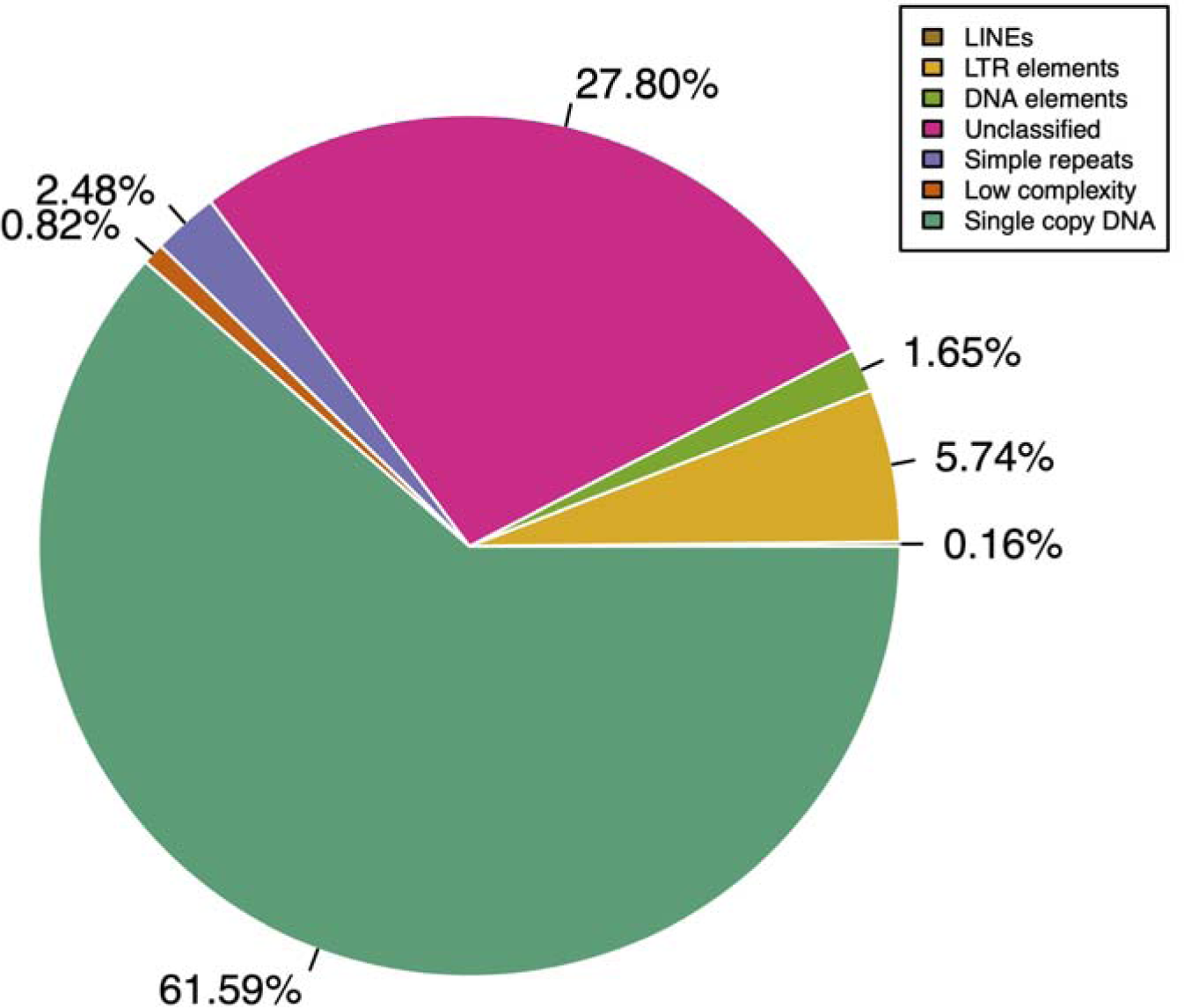
Repeat content classification using RepeatModeler2 and RepeatMasker. The proportion per category of repetitive element found across the genome. Categories include long interspersed elements LINEs (brown), LTR elements (yellow), DNA elements transposons (light green), simple repeats (purple), low complexity (orange) and unclassified repeats (pink).

**Table 5.**
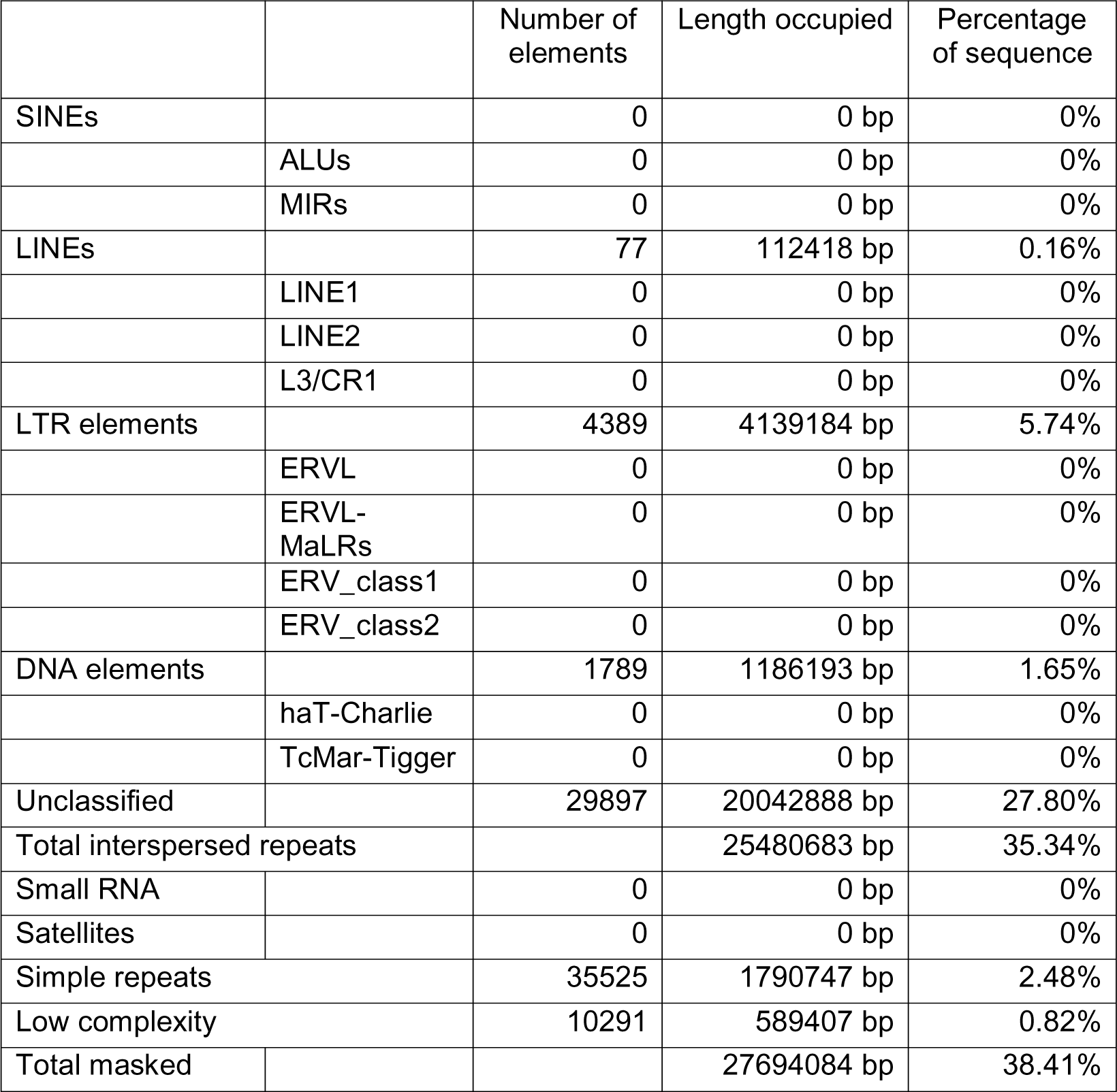
Repetitive elements modeled de novo with RepeatModeler2 (class and content).

### Annotation

We annotated the *X. peckii* genome assembly using the BRAKER3^49^ pipeline. First we used RepeatMasker v.4.0.734^48^ to soft mask the genome (--xsmall) using the repeat library we generated de novo with RepeatModeler2^47^. Next we used the software STAR (Spiced Transcripts Alignment to a Reference) v2.7.1^50^ to map RNA-seq reads (male; whole body) to the genome. Then we used the mapped reads (--bam) as evidence input for BRAKER3^49^ to generate ab initio gene predictions which resulted in 8759 predicted genes. Additionally, we did a homology-based gene prediction with the tool GeMoMa v1.9^51^. We implemented the annotated proteins from the published genome of *Drosophila melanogaster* as reference input with the following command: java -Xmx126976m -Xms126976m -jar GeMoMa-1.9.jar CLI GeMoMaPipeline AnnotationFinalizer.r=NO o=true t=./UROC_Xpeckii_1.1.fasta s=own i=Dmel a=dmel-all-r6.45.problem.free.subset.gff g=dmel-all-chromosome-r6.45.fasta r=MAPPED ERE.s=FR_FIRST_STRAND ERE.m=C641Aligned.sortedByCoord.out.bam threads=16 outdir=annotation_run1. The combination of homology-based and ab-initio gene predictions resulted in 8783 genes.

## Data Records

This Whole Genome PacBio HiFi assembly project has been deposited at DDBJ/ENA/GenBank under the accession JAWUEG000000000. The raw genomic and transcriptomic Illumina PE sequencing reads were deposited at the NCBI Sequence Read Archive with the Bio project accession’s number PRJNA1032868 BioSample accession numbers are SAMN39879530 (Male WGS), SAMN39879531 (females WGS) and SAMN39879532 (Male RNA seq). The GFF3 files with ab initio gene predictions from BRAKER and the homology-based gene predictions from GeMoMa are available in FigShare repository.

## Technical Validation

### GenomeScope2 size estimation validation and comparative analysis with other *Xenos* species

We used the short 150bp Illumina PE reads from both male and female *X. peckii* samples to verify the accuracy of our GenomeScope2 estimations from the raw PacBio HiFi reads. Next, computed the histogram of k-mer frequencies with Jellyfish v2.3.0^29^ (-C -m 21; histo –h 1,000,000), and ran the output through the online web tool of GenomeScope2 (K-mer length = 21, ploidy = 2 and max K-mer coverage = 1,000,000)^30^. The estimated genome size using male and female high-coverage short Illumina reads suggests that the genome size ranges between 67,923,714 bp for the male and 75,093,981 bp for the female (Table 3; Fig S4). Moreover, the analysis from both sets of reads estimate that the repetitive content of the genome is around 43.7% and 41.4% for the male and female samples, respectively.

Additionally, to further validate our results and place them in a comparative framework, we performed the same GenomeScope2 estimations with the only two *Xenos* species for which whole genome shotgun sequence data is available: *X. vesparum* and *X.cf. mouton* (recently reported to be the same sspecies as *X. yangi*) (150bp Illumina PE reads; NCBI Accession number SAMN03323551 and PRJNA681068 respectively). The genome size of *X. vesparum* was previously calculated using flow cytometry data and is reported to be ∼ 133Mb^6^. Our GenomeScope estimations are consistent with this previously reported size for *X. vesparum,* with a slightly smaller genome size of ∼ 120.8 Mb. Similar to our *X. peckii* assembly, the *X. vesparum* data showed small discrepancies (∼10Mb) between the GenomeScope2 estimations and the genome size calculated using flow cytometry. These results indicate that our estimations closely reflect the genome size of the 3 *X. peckii* samples that we analyzed. Moreover, our results reveal a significant amount of variation in genome size in *Xenos*, ranging from 58,068,699 bp in *X. yangi* to 120,869,754 bp in *X. vesparum* (Table 3). Importantly, the genome size estimations for our male *X. peckii* reads are ∼10 Mb smaller than the female reads which is consistent with differentiated XY chromosomes. If the X chromosome has slightly higher repetitive content, then we would expect the females to have a larger genome size. However, the repetitive content estimated for both male and female short reads was similar (Table 3). Further studies will confirm if the difference in genome size estimations is due to sequencing or assembly artifacts or due to real biological differences in the genomes of males and females.

### Genome assembly metrics with different software

We ran four different assembly software (i.e. Hifiasm^31^, Improved Phased Assembler IPA^32^, Hicanu^33^, Flye^34^) to assess which one produced the highest quality genome. We evaluated the quality for all assemblies using QUAST v.5.0.2 ^38^ (Supplementary Table 1; Supplementary Figure 1) and BUSCO v.5.2.2 ^39^ with the insecta_odb data base (Supplementary Figure 2). Hifiasm produced the overall most contiguous and complete genome assembly, with the lowest rates of duplications reported by BUSCO. Both Flye and Hicanu were inefficient in resolving the assembly, which resulted in large and fragmented assemblies with high duplication according to the BUSCO scores (Supplementary Figure 2). Moreover, we used a *Jupiter plot* ^35^ (-n=50000, ng=75) to compare the contiguity and completeness of the assemblies produced by different software. We confirmed that the assembly produced by Hifiasm is more contiguous and complete than the one produced by IPA which was the second-best ranked assembly of our analysis (Supplementary Figure 7).

## Code Availability

No specific code or script was used in this study. Data processing was executed following the documentation of the corresponding software described in the methods. Software programs with no parameters associated were used with the default settings.

## Acknowledgements

This work was supported by funding provided by the University of Rochester to M.I.C. and F.M.K.U. Special thanks to J. Albert C. Uy, Faye Romero, Christine Muirhead, Jack Werren and Felix Beaudry for invaluable advice. Juan Martín Ferro and Emiliano Martí provided crucial advice for assembling the mitogenome and code to run the genome annotation. Amanda Larracuente, and members of the TropBioLab at the University of Rochester provided feedback which significantly improved this manuscript. Erin Bernberg, Olga Shevchenko and Brewster Kingham at the University of Delaware sequencing facility provided thoughtful logistical support. We thank Sloan Tomlinson and Adam Fenster for insect photographs, and James Brophy at Robert H. Truman Park for field support. Samples were collected under permit OPHRP FL01 issued by the New York Office of State Parks.

## Author contributions

M.I.C. and F.M.K.U. designed the study. F.M.K.U collected the samples and performed DNA and RNA extractions. M.I.C. performed the analyses. X.Y. provided valuable code and input to perform the genome assembly. M.I.C and F.M.K.U. wrote the first version of the manuscript. All authors read, revised and approved the final manuscript.

## Competing interests

The authors declare no competing interests.

## Notes

### Competing Interest Statement

The authors have declared no competing interest.

