## Supplementary material for "First genome assembly of the order Strepsiptera using PacBio HiFi reads reveals a miniature genome"

Table of contents

Supplementary Figure 1 – Page 2

Supplementary Figure 2 – Page 3

Supplementary Figure 3 – Page 4

Supplementary Figure 4 – Page 5

Supplementary Figure 5 – Page 6

Supplementary Figure 6 – Page 7

Supplementary Figure 7 – Page 8

Supplementary Table 1 – Page 9

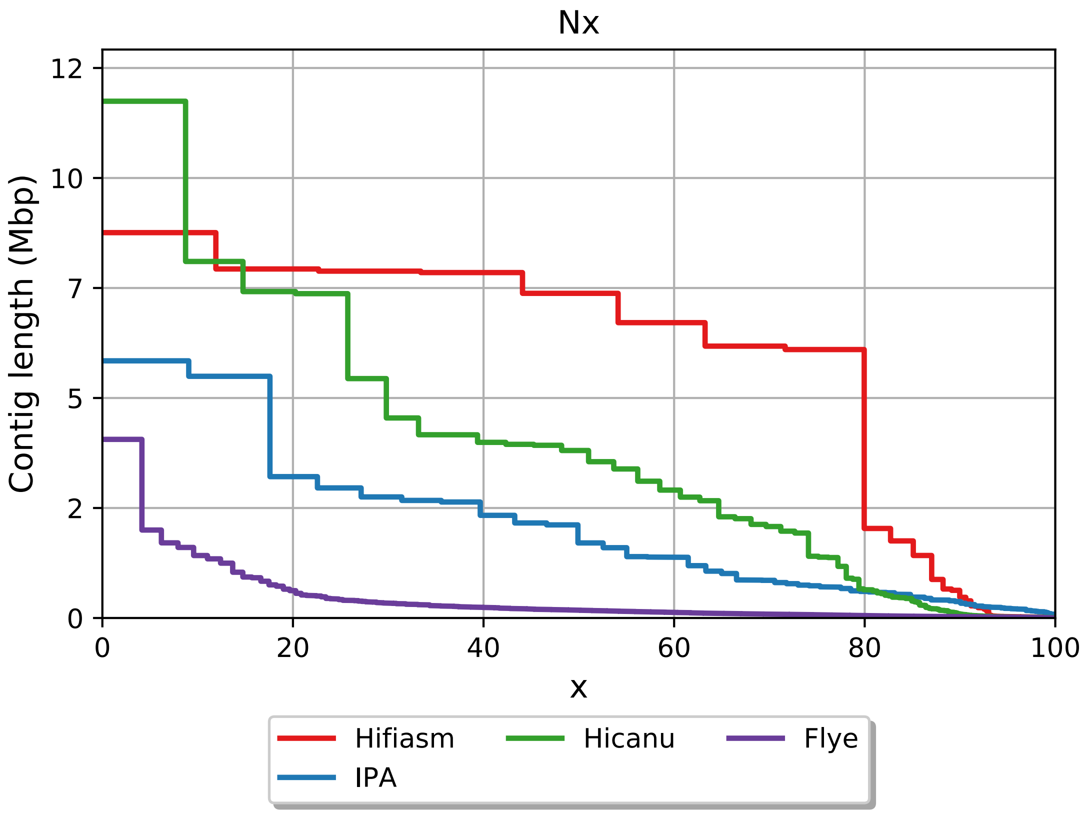


**Figure S1.** QUAST comparison of assembly contig length produced by different software.

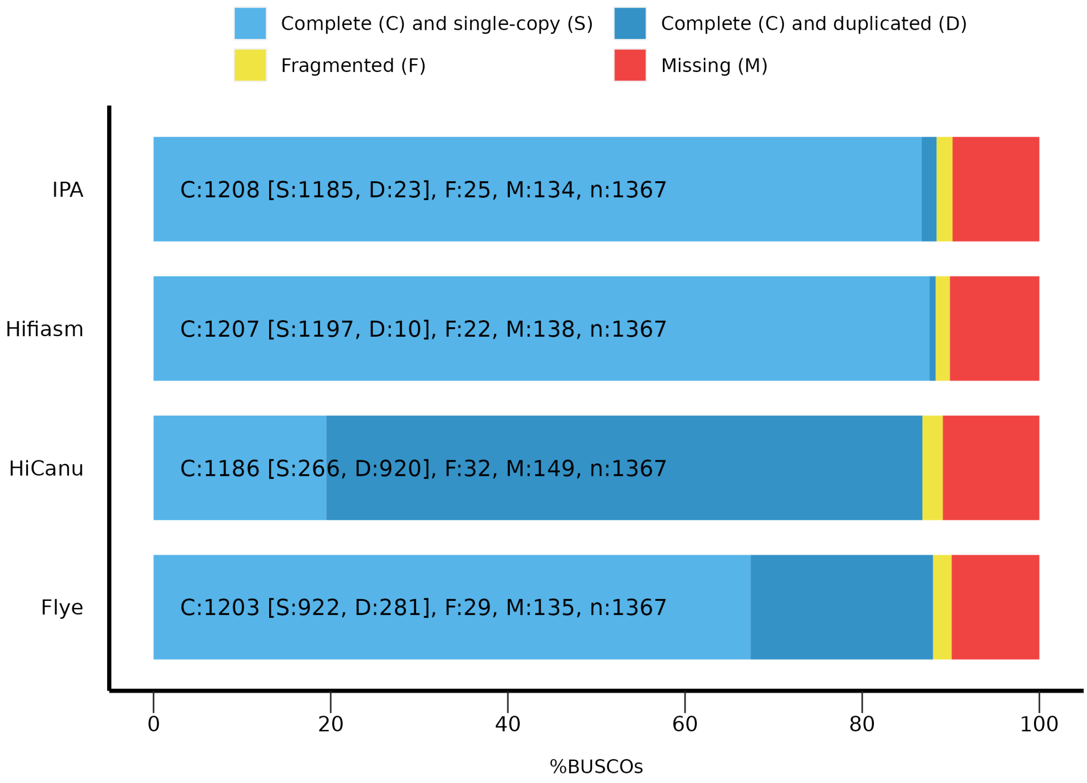


**Figure S2. BUSCO scores for assemblies produced by different software.**

Assembly with Hifiasm shows the highest completeness score with lowest duplication rates. Approximately 10% of the genes in the BUSCO database are missing from the assembly for all independent assemblers.

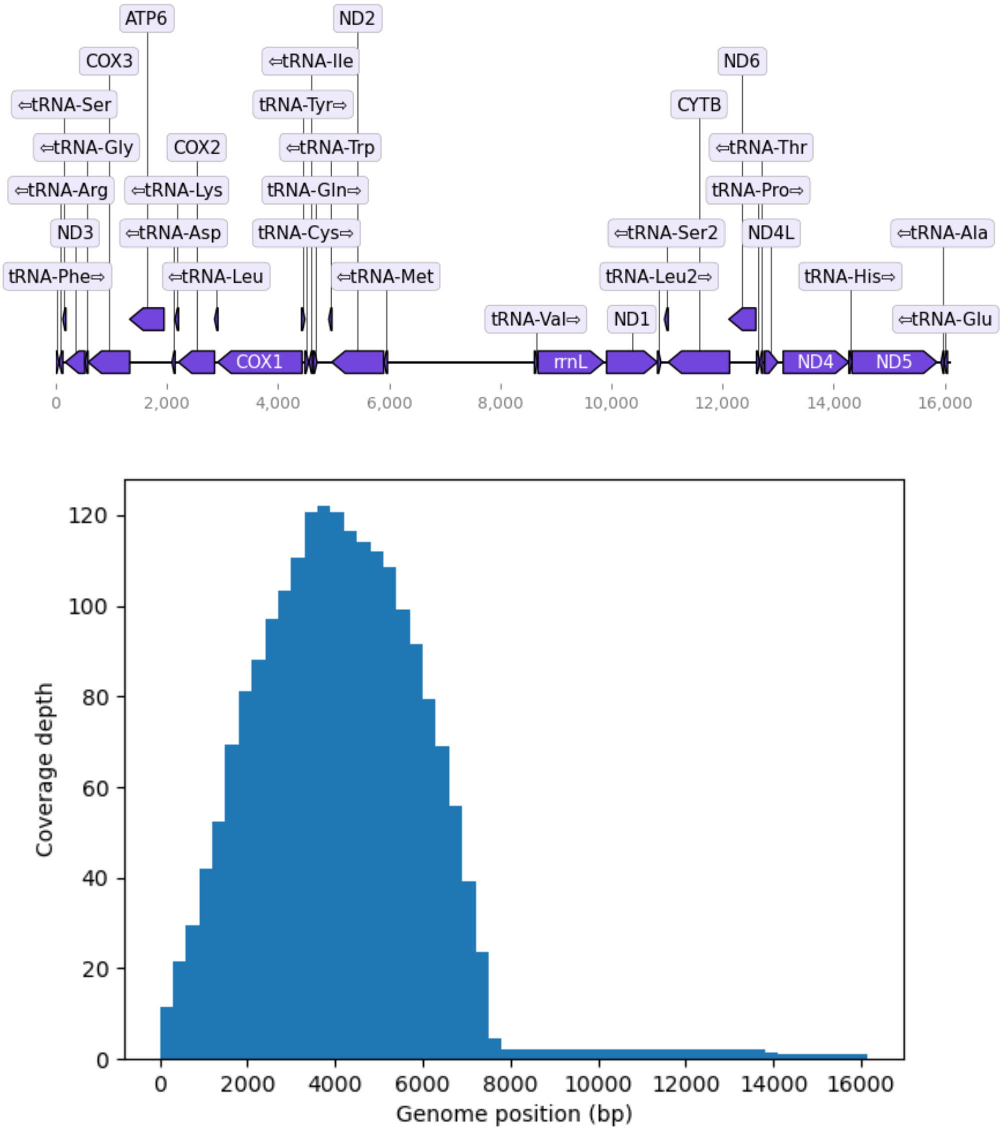


**Figure S3**. Final mitogenome annotation (top) and coverage along the sequence (bottom).

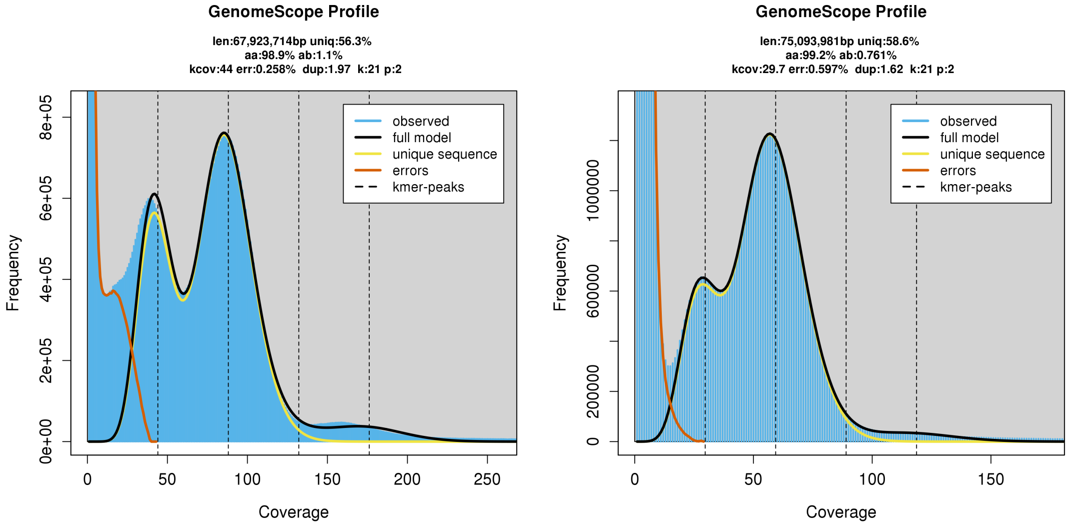


**Figure S4.** Genomescope2 assessment using male (left) and female (right) short Illumina PE reads. Overview of the k-mer frequency distribution for the *X. peckii* genome. X axis – k-mer depth, Y axis = k-mer frequency for a given depth.

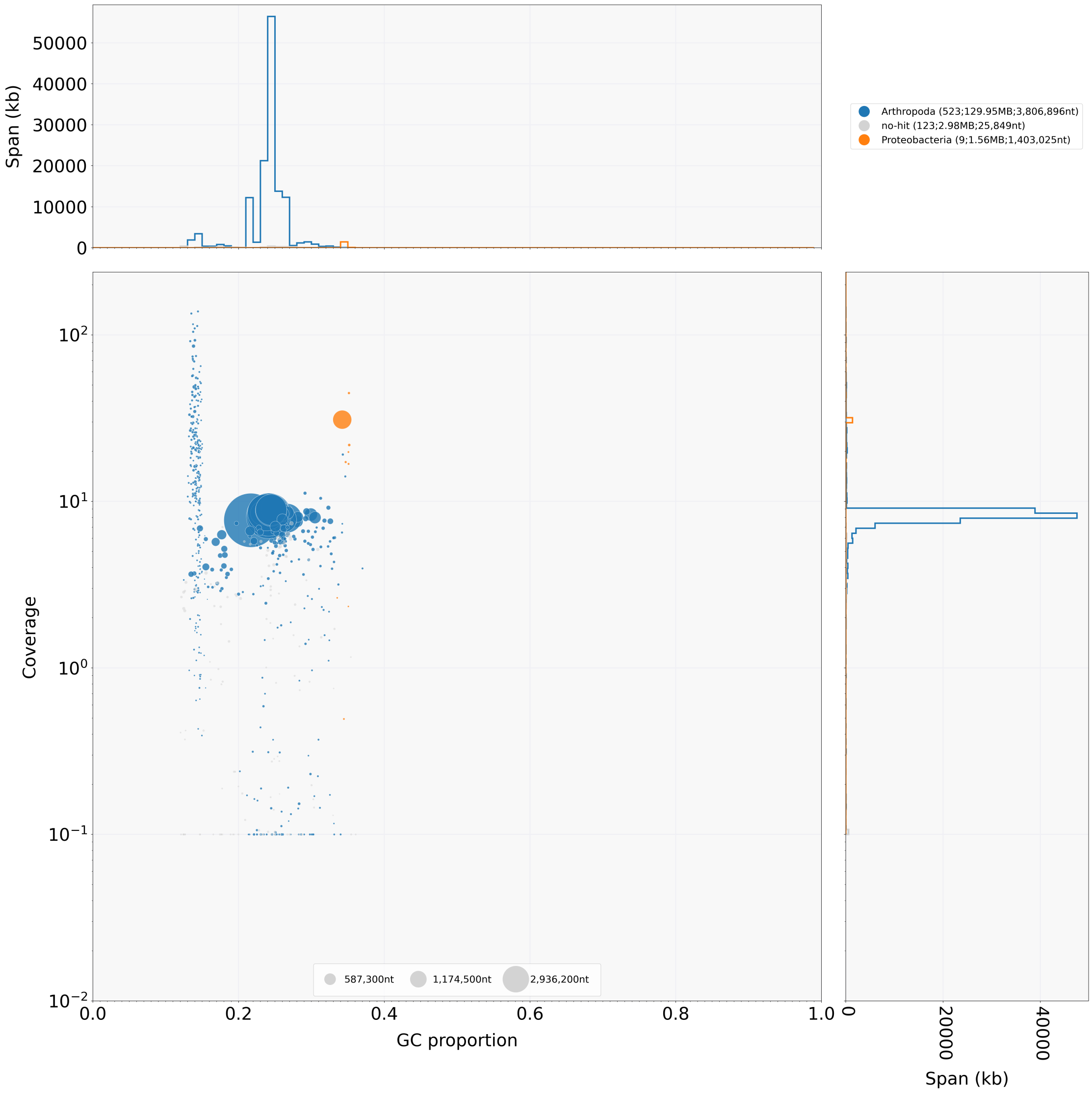


**Figure S5.** Blobtools scan for contamination results for the sequenced male *X. peckii*.

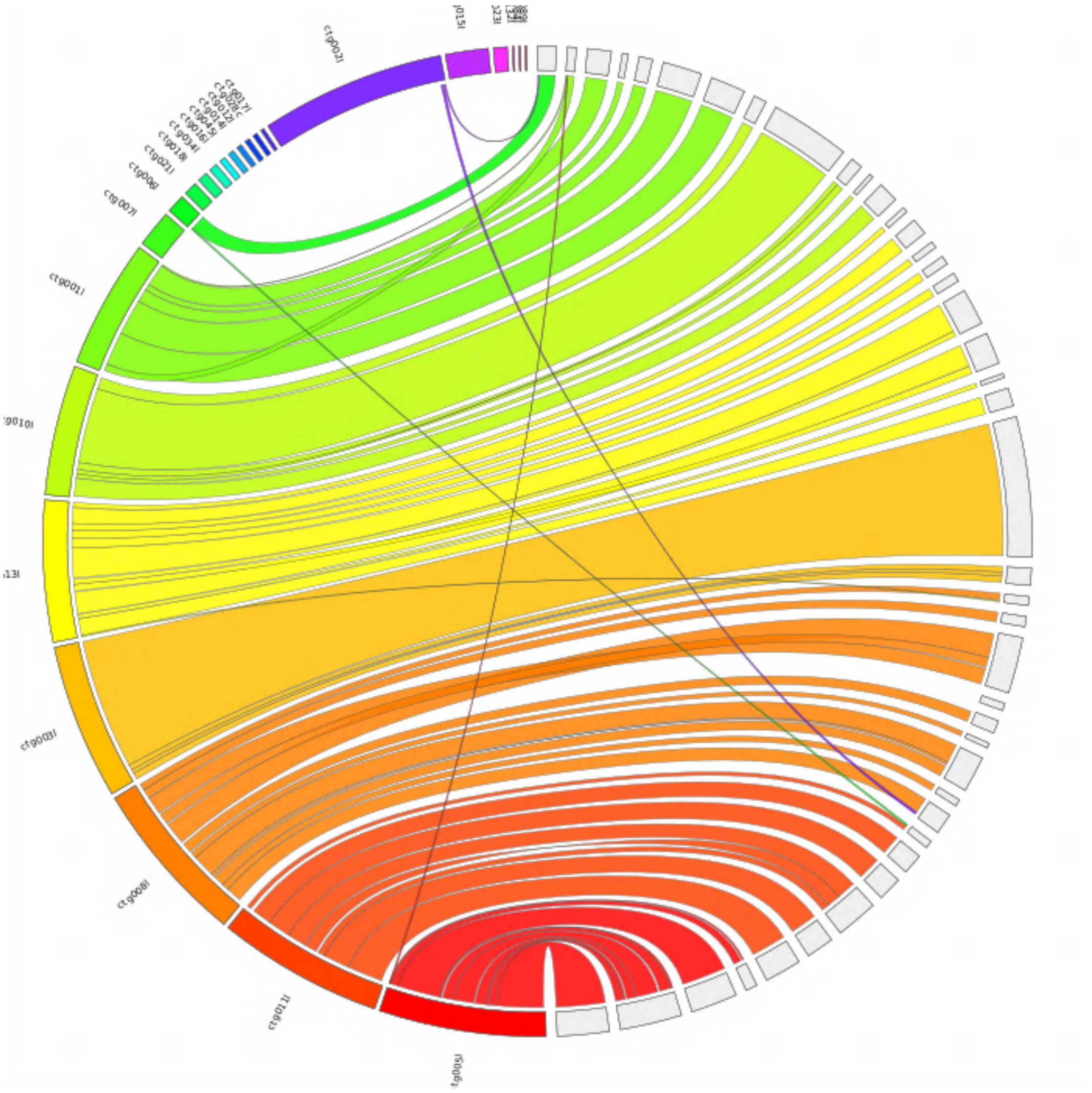

**Figure S6. Jupiter synteny plot of Hifiasm primary vs. alternate assemblies.** Synteny between the primary (left) and alternate (right) phased assemblies produced by Hifiasm. The primary assembly is more contiguous and includes two large contigs that show coverage patterns consistent with sex chromosomes, compared to the alternate assemblies.

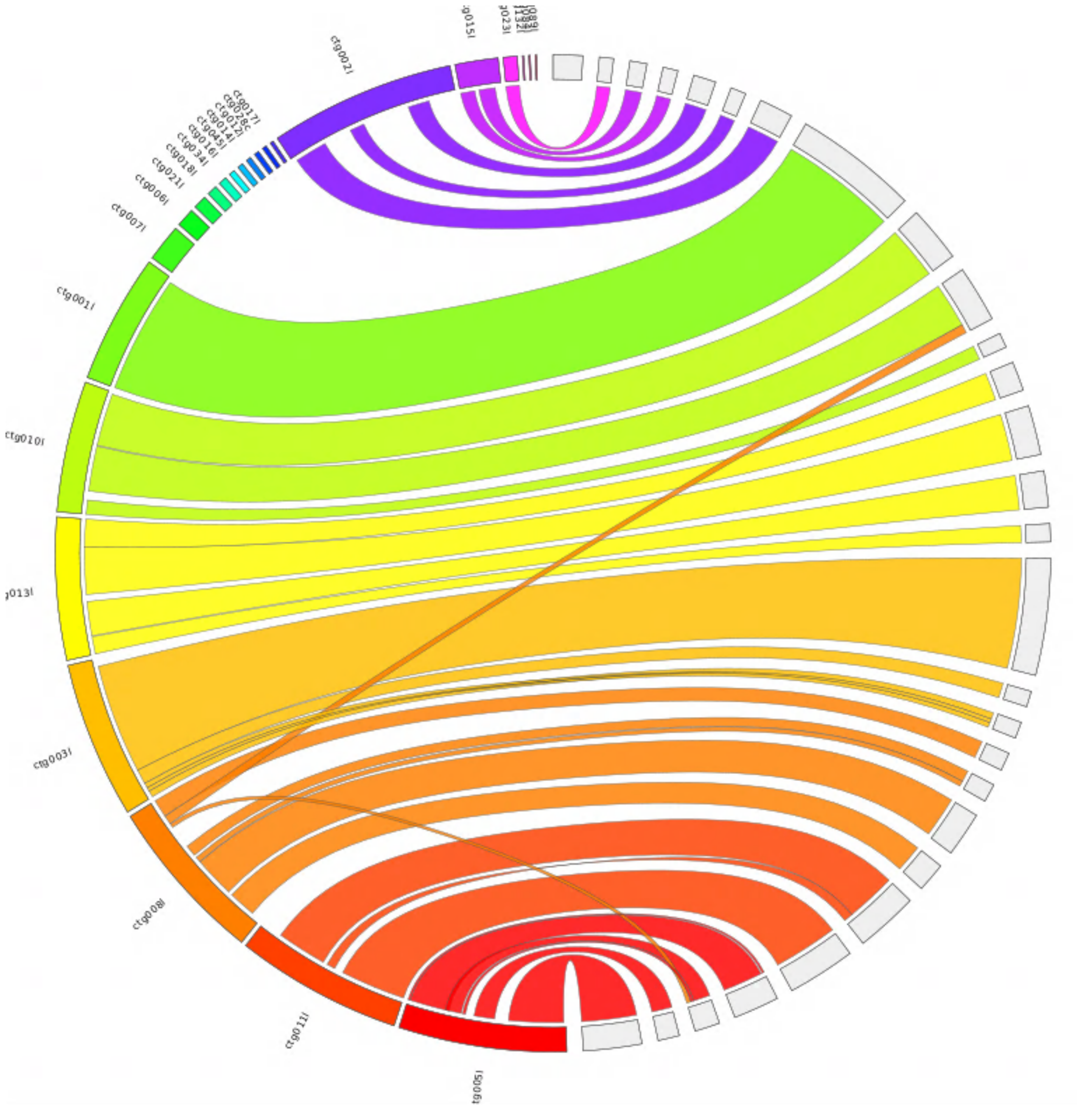


**Figure S7. Jupiter synteny plot of assemblies with Hifiasm vs. IPA.**

Synteny between assemblies produced with different methods, Hifiasm(left) IPA (right). Assembly produced by Hifiasm is more contiguous (higher N50) but shows relatively consistent synteny between both assemblies.

**Table S1. Assembly statistics using four different assembly software.**

| | Hifiasm | IPA | Hicanu | Flye |
|---|---|---|---|---|
| Total length | 73,541,611 | 64,473,870 | 123,502,800 | 97,837,690 |
| # Contigs | 229 | 77 | 655 | 1299 |
| Largest contig | 8,758,233 | 5,844,905 | 11,744,670 | 4,059,769 |
| GC % | 23.61 | 24.59 | 24.05 | 24.05 |
| N50 | 7,378,620 | 1,706,016 | 3,806,896 | 177,262 |
| N75 | 6,101,180 | 732,661 | 1,403,025 | 72,594 |
| L50 | 5 | 11 | 12 | 121 |
| L75 | 8 | 25 | 25 | 341 |
